# Improvement of thermostability and activity of PET-degrading enzyme Cut190 towards a detailed understanding and application of the enzymatic reaction mechanism

**DOI:** 10.1101/2023.02.26.529345

**Authors:** Nobutaka Numoto, Narutoshi Kamiya, Masayuki Oda

**Affiliations:** Medical Research Institute, Tokyo Medical and Dental University, Tokyo 113-8510, Japan; Graduate School of Information Science, University of Hyogo, Hyogo 650-0047, Japan; Graduate School of Life and Environmental Sciences, Kyoto Prefectural University, Kyoto 606-8522, Japan

## Abstract

Enzymes capable of hydrolyzing polyethylene terephthalate (PET) and other plastics are attractive catalysts for application to the recycling of plastic waste due to their generally low environmental impact. Cut190 is a cutinase from a thermophilic actinomycete and shows PET-degrading activity and high thermal stability. We developed a series of Cut190 mutants exhibiting further improvements in thermal stability and activity, and showed that the unique stabilization and activation mechanism was dependent on Ca^2+^ ions. Two of these mutants, Cut190** and Cut190*SS, differed from the previous mutant Cut190* by deletion of the three C-terminal residues and introduction of five substitutions, including two cysteines forming a disulfide-bond, respectively. These mutants exhibit higher thermal stability and activity, which are often mutually exclusive characteristics. Crystallographic studies of these mutants and their inactivated derivatives demonstrated that they could have a novel ejecting form that would be responsible for releasing products. We also determined the crystal structures of ligand-bound complexes, which revealed the molecular mechanisms of the aromatic-ring recognition and the tetrahedral intermediate during the substrate cleaving, although the ligands had no aromatic ring but a cyclic group. This structural information provides insights into the mechanism of the Ca^2+^ -dependent PET-cleaving activity of Cut190 and provides a useful basis for further mutant design and computational studies.

## Introduction

Enzymatic recycling of plastic wastes such as polyethylene terephthalate (PET) by naturally occurring enzymes would be an ideal solution to their mounting environmental ubiquity because such enzymes have low environmental impact. Several enzymes capable of hydrolyzing PET have been reported (*1*). Most of them are cutinases (EC 3.1.1.74), which were originally identified as enzymes degrading cutin (Figure 1a), the main component of the cuticle layer of leaves. Cutin is an aliphatic polyester, whereas PET is a polyester containing aromatic rings. The molecular mechanism by which cutinase recognizes and degrades polymers with aromatic rings is the key to the development of applications for this enzyme, and is also of interest in protein science. To improve the thermal stability of the enzyme is particularly important for the practical application of enzymatic degradation of PET, because an enzymatic reaction at 70°C or higher, which is above the glass transition temperature of PET, is necessary to efficiently access and degrade the polymer chains of PET.

**Figure 1.**
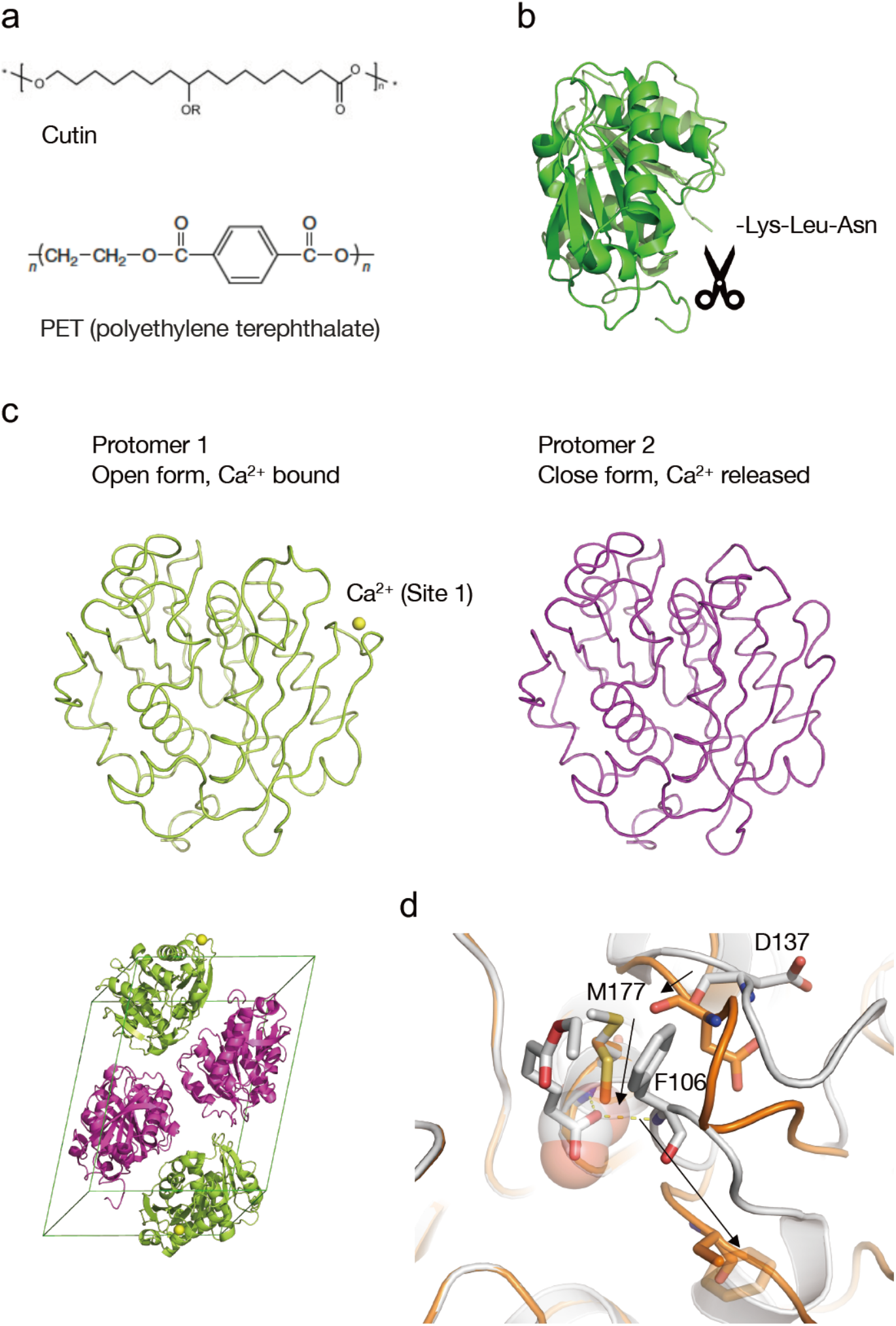
Structure of Cut190**. (a) Substrates of cutinase. (b) Ribbon diagram of Cut190 and the C-terminal three residues that were deleted in Cut190**. (c) Protomers in the crystal of Cut190**. The bound Ca^2+^ ion at Site 1 is drawn as a yellow sphere in protomer 1, whereas no Ca^2+^ ion is observed in protomer 2 (magenta). The lower left panel shows the protomers in the unit cell (twice the asymmetric unit). (d) Closeup view around the active site. The ejecting form of Cut190**S176A (orange) is superposed onto the open form of Cut190* complexed with monoethyl adipate (white, 5ZRS). Remarkable structural changes are indicated as black arrows. The side chain of Met177 in the ejecting form causes the steric hindrance to the bound ligand at the active site. The aromatic ring of F106 in the ejecting form is disordered (drawn as a transparent stick) in the crystal structure.

A cutinase from a thermophilic actinomycete *Saccharomonospora viridis* AHK190, known as Cut190, has a high thermal stability with a *T*_m_ value above 55°C in the wild type (*2*). Cut190 would therefore be promising as a target enzyme for modification to improve the thermal stability and enzymatic activity. We previously reported the biochemical properties of Cut190, and we clarified that its thermal stability and activity are greatly affected by divalent cations, with Ca^2+^ ions in particular improving both the thermal stability and activity (*2-4*). Mutational and structural studies demonstrated that there are three major Ca^2+^ binding sites in the enzyme, with two (Sites 2 and 3) being primarily responsible for the thermal stability and the other (Site 1) being responsible for the activity (*5, 6*). Interestingly, large structural changes around Site 1 associated with the structural changes at the active site regulate the enzymatic activity via an allosteric mechanism. Our structural studies, including an analysis of substrate-bound forms (*3, 5*), showed that there are at least three forms—a closed, an open, and an engaged form—with calcium binding and dissociation during the reaction cycle. Collectively, these results provide insights into the molecular mechanism underlying the structural changes coupled with the dynamic binding and dissociation of Ca^2+^ ions during the enzymatic reaction cycle (*5*). In parallel with these structural studies, we developed several mutants with improved thermostability and activity based on the structural information (*2, 6*). Recently, we developed further-improved mutants (*6, 7*) that show even higher thermal stability and activity, two characteristics that are often mutually exclusive.

The successful development of engineered enzymes designed based on a machine learning algorithm has been reported recently (*8*). However, rational mutant design based on detailed structural information, including the structural information of substrate complexes, is required because Cut190 is a unique calcium-dependent enzyme among enzymes that exhibit PET-degradation activity. In the present study, we determined the crystal structures of improved mutants of Cut190, in ligand-bound forms. Compared with the previously published structural bases of substrate/ligand-bound forms (*5, 7, 9*), these structures provide more detailed insights into the molecular mechanism of the enzymatic activity and useful structural information for the rational design of Cut190 mutants that exhibit further improvement of PET-degrading activity.

## C-terminal deletion improved both thermal stability and activity

The first successful mutant having both higher thermal stability and higher activity than the wild type was Cut190 (S226P/R228S), which we named Cut190* (*10*). Recently, we developed a further-improved derivative of Cut190*, named Cut190**(*7*), in which the C-terminal three residues of Lys-Leu-Asn were deleted (Figure 1b). Both the thermal stability and enzymatic activity were improved over those of Cut190* (Table 1).

**Table 1.** Melting temperatures of Cut190 mutants with 2.5 mM Ca^2+^ ions.

| | $T_m$ (°C) | $K_{cat}/K_m$ (mM<sup>-1</sup>s<sup>-1</sup>) |
| --- | --- | --- |
| Cut190* | 68.4 | 308 |
| Cut190** | 72.7 | 357 |
| Cut190*SS | 83.3 | 327 |

Data were taken from Oda *et al.*, 2018 (6), Senga *et al.*, 2021 (7), and Emori *et al.*, 2021 (9).

To elucidate the molecular mechanism underlying the improved enzymatic functions, we determined the crystal structure of Cut190** (*7*). To our surprise, one protomer showed an open form with a Ca^2+^ ion at Site 1, while the other had a closed form without a Ca^2+^ ion in the crystal that contained two molecules in the asymmetric unit (Figure 1c). Since the crystallization condition contained 5 mM of Ca^2+^ ions, these facts strongly suggest that the Ca^2+^ ion binding at Site 1 would be transient, and the ions could be released with large conformational changes around Site 1. This is supported by our previous Ca^2+^ binding analysis using 3D Reference Interaction Site Model (RISM) calculations, in which Site 1 is ranked as having relatively low affinity (*5*). Focusing on the C-terminal structure, we found that a salt bridge is formed between the carboxyl group of C-terminus Y304 and the side chain of R274 in both the open and closed forms. However, we also found that the Δ*H*_cal_ values of the C-terminal Lys-Leu-Asn deletion mutant at the respective Ca^2+^ ion concentrations were similar or lower than those of the C-terminal Lys-Leu-Asn-containing mutant in our thermodynamic analysis of differential scanning calorimetry (DSC) data, indicating that the increased thermal stability of Cut190** is due to entropy effects (*7, 10*). We also determined the crystal structure of the Cut190**S176A mutant (*7*), which was inactivated through substitution of the active Ser176 to Ala. The structure revealed a novel form that is different from any of the known structures of closed, open, or engaged forms (*3, 5*). The most significant structural change occurs at the β3-α2 loop including Phe106 (Figure 1d), whose main chain nitrogen is responsible for the oxyanion hole. The position of the Cα atom of Phe106 moves 6.6 Å, resulting in disruption of the oxyanion hole, which is indispensable for stabilization of the reactive intermediate state of the substrate. Moreover, the rare side chain rotamer of Met177 will cause a large steric hindrance and prevent substrate binding. These structural changes indicate that the structure of Cut190**S176A is in an inactivated, ligand-incompatible state and suggests that the changes promote dissociation of the bound substrate or product. The structural changes around Site 1 are also remarkable, because the conformation is different from any reported structures. No electron density of the bound Ca^2+^ ion is observed, even though the conformation of the β1-β2 loop responsible for forming Site 1 is similar to that of the Ca^2+^ bound open form. Because the Ca^2+^ binding site at Site 1 is composed of the main chains of the β1-β2 loop, even small structural changes will have a large impact on the manner of Ca^2+^ ion coordination. Thus, we designated the structure of Cut190**S176A as an ejecting form that exists in a product-releasing state between just after the enzymatic reaction and just before the binding of the next substrate.

Integrating the above results, we updated the reaction cycle of Cut190 based on the crystal structures (Figure 2). The closed, substrate inaccessible form is activated to the open, substrate-waiting form induced by Ca^2+^ binding at Site 1 (I to II). A substrate can access the active site, but the open form would not be suitable to stably accommodate the substrate (*5*). Then, structural change to the engaged form (III) accompanying Ca^2+^ ion release is required to hold the substrate tightly and initiate cleavage at the active site. Once the enzymatic reaction is complete, the active site should be open for the rapid exchange between the product and new substrate (IV). In this step, the Ca^2+^ ions will be bound at Site 1 again. Previous molecular dynamics simulations have demonstrated that ligand molecules would be rapidly dissociated in the open form. Nevertheless, there is still a possibility that the reaction would stall at the open form and the product could not dissociate in the active site, or that the product would revert to the engaged form, in which the ligand would be tightly accommodated again. The ejecting form forces the product to dissociate from the active site (V) and allows the substrate exchange to start the next reaction cycle.

**Figure 2.**
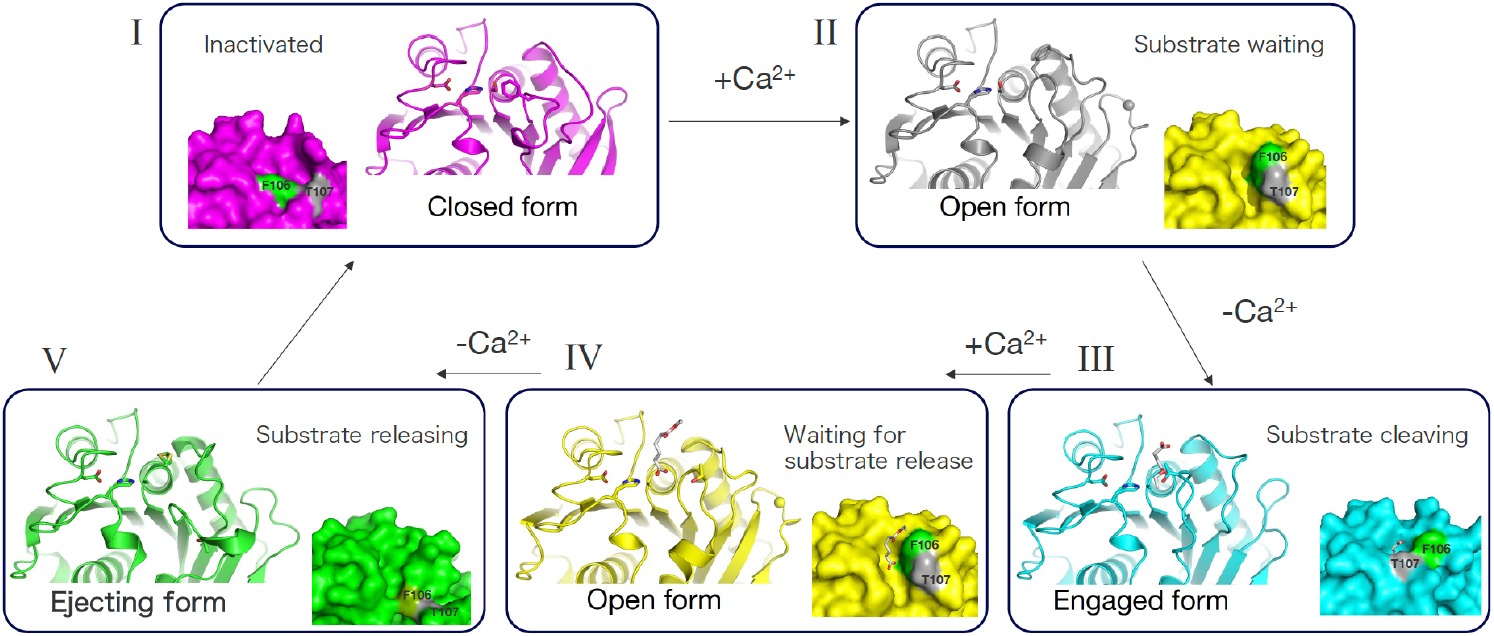
Schematic diagram of the enzymatic reaction cycle of Cut190. A novel ejecting form is integrated. Ca^2+^ ions are dynamically bound and dissociate in the cycle to promote structural changes.

## Introducing a disulfide bond significantly improved the thermal stability

Our continuous search for mutants led us to the mutant with the highest thermal stability above 80°C (Table 1). This mutant, named Cut190*SS(*9*), was based on Cut190* but contained five additional substitutions, i.e., Q138A/D250C-E296C/Q123H/N202H (*6*).

Notably, Cut190*SS exhibited both a significant improvement in thermal stability and high activity compared to Cut190*. In particular, the introduction of two cysteines (D250C and E296C) was intended to form a disulfide bond at the Ca^2+^ binding site (Site 2), which disulfide bond made the primary contribution to the thermal stability of Cut190* (*6*). Thereby we expected to obtain more effective thermostability than Ca^2+^ dependent stability. We then determined the crystal structure of Cut190*SS (*9*) to elucidate the molecular mechanism of the improved functions of Cut190*SS. The structure was determined as a closed form (Figure 3). No significant structural changes were induced by introducing the five substitutions. As expected, two Cys residues (D250C and E296C) formed a disulfide bond and a Ca^2+^ ion was no longer bound at this site. We also determined the structure of an inactivated derivative of Cut190*SS_S176A (*9*), whose active Ser176 is replaced with an Ala. The structure shows the open form, and only Site 1 binds a Ca^2+^ ion. There are no significant changes in the overall structure compared to the previously determined open form of Cut190*, except that a disulfide bond is formed at D250C and E296C. Therefore, these structures confirm that five substitutions containing a disulfide bond formation did not induce any significant changes to the overall structure, and did not affect the ability to exchange the closed and open forms, which is crucial to regulate the enzymatic activity. The substitution of Q138A contributed to the maintenance of higher activity because Gln138 is located near the substrate-binding site, as demonstrated by the activity measurements of the single mutation (*9*). The smaller sidechain of Ala seems to be more suitable to easy access of the substrate to the active site.

**Figure 3.**
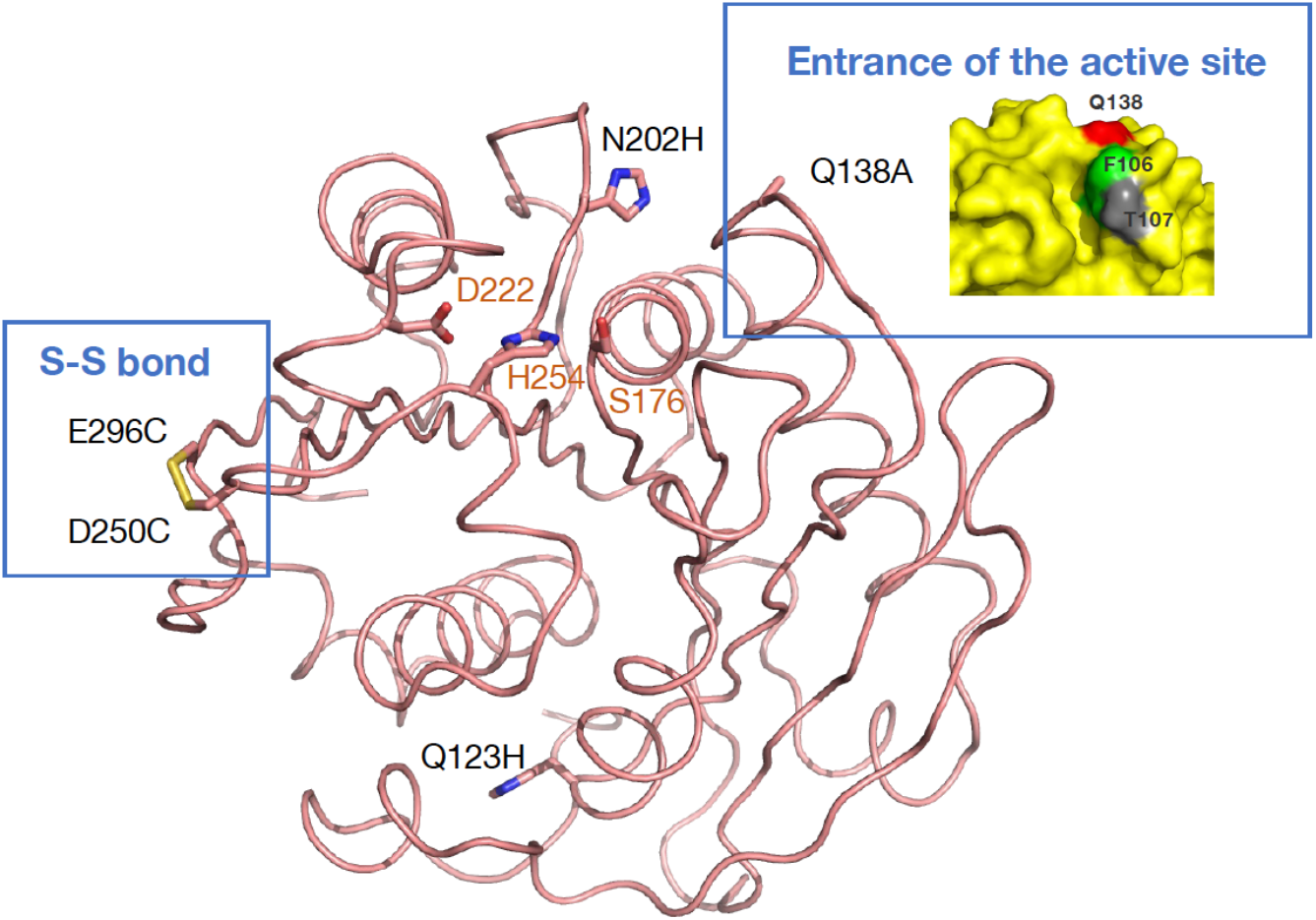
Crystal structure of Cut190*SS. Five substitutions introduced in Cut190* and the residues of S176, H254, and D222 that compose the active site and so-called catalytic triad are indicated and shown as stick models. Substitution of Q138A is located at the entrance of the active site (inset panel at upper right) and would affect the substrate binding. F106 and T107 function as a lid for the active site.

## Determination of ligand-complex structures

To develop a more improved enzyme, it is obviously essential to obtain the structural bases of the complex of the enzyme and substrates for the rational design of mutants. We have previously reported the crystal structures of the substrate-bound forms of the Cut190* S176A mutant (*5*). We used small aliphatic esters, monoethyl succinate and monoethyl adipate, for the determination of the crystal structures. Though the substrates contain no aromatic ring characteristic of PET, the complexed structures have provided useful insights into the molecular mechanism and the enzymatic reaction cycle with remarkable structural changes. Continuing attempts to analyze complex structures with PET-like substrates containing aromatic rings are ongoing. We recently succeeded in determining the structures of complexes with molecules having a ring structure, although the ring in these structures is different from an aromatic ring. One of these was a complex of Cut190*SS_S176A and dioxane (Figure 4a) as reported previously (*9*), and another was a complex of Cut190*SS_S176A and 2-(N-morpholino)ethanesulfonic acid (MES). In addition, we also determined the structure of a complex of Cut190*SS_S176A and a sulfate ion at the oxyanion hole. These structures provide important insights into the manner of accommodation of the tetrahedral intermediate for the PET cleaving. diffraction experiments were performed at BL-17A at KEK-PF using an automatic data-collection system (*11*), and the data were automatically processed by PReMo (*12*). Manual data processing using XDS (*13*) and truncation using the CCP4 program suite (*14*) were performed as necessary. Phases were determined by the molecular replacement method using PHASER (*15*). Several cycles of manual model rebuilding and refinement were performed by using Coot (*16*) and PHENIX (*17*), respectively.

**Figure 4.**
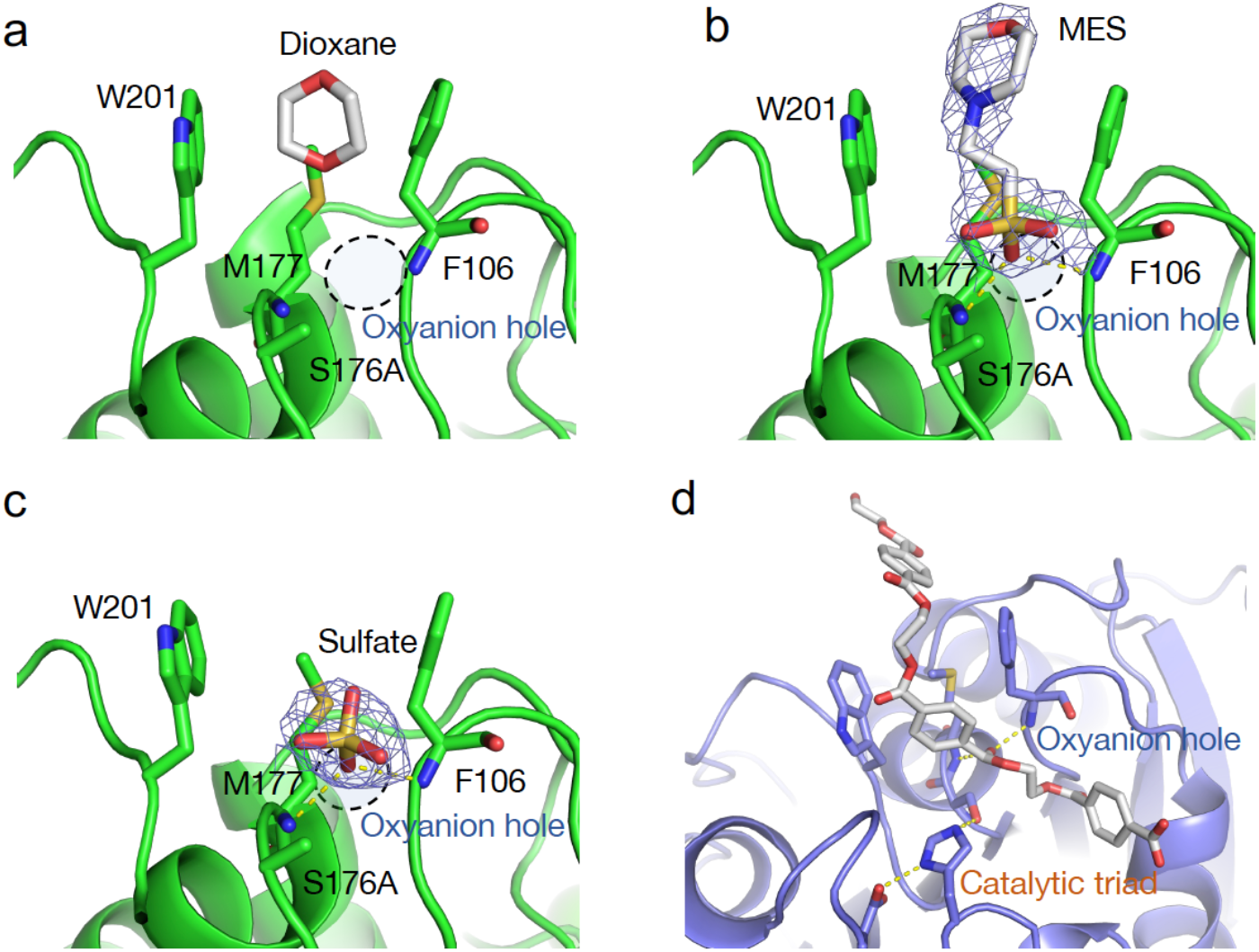
Ligand-bound structures of Cut190*SS_S176A. (a-c) The dioxane-, MES-, and sulfate-bound forms, respectively. The active Ser is mutated to Ala. Oxyanion holes composed of the main chains of F106 and M177 are indicated as dashed circles. For the MES- and sulfate-bound structures in this study, electron density (2F_o_-F_c_, contoured at 1σ level) around the bound ligands is shown as blue mesh. Hydrogen bonds between the ligands and oxyanion hole are drawn as yellow dashed lines. (d) Model structure of PET-3mer (white stick)-bound Cut190. Interactions at the catalytic triad and oxyanion hole are represented as yellow dashed lines.

The structure of the MES complex (Figure 4b) was determined at 2.6 Å resolution (Table 2). Strong residual electron density was observed at the active site, but when we assigned the synthetic substrate, the molecule was a little smaller than the residual density. After careful consideration of the possible molecules included under the crystallization condition, we concluded that the bound ligand is MES with a ring-shaped morpholine group and a sulfo group. The morpholine group is accommodated in aromatic residues of Phe106 and Trp201. This manner of accommodation by a ring-shaped molecule is almost the same as observed in the structure of Cut190*SS complexed with dioxane (*9*). Moreover, the structures of leaf-branch compost cutinase (LCC) complexed with dioxane have also been reported (*18*), and the dioxane-binding modes are in good agreement between Cut190 and LCC. Therefore, this site would prefer to bind hydrophobic ring-shaped molecules. These facts strongly suggest that the aromatic ring moiety of bound PET will be located at the same dioxane-binding site between Phe106 and Trp201. On the other hand, a sulfo group is recognized in the oxyanion hole composed of the main chain amines of Phe106 and Met177 through the hydrogen bonds. The side chain of His254 of the canonical catalytic triad also forms a hydrogen bond with the sulfo group. These facts strongly suggest that the bound MES molecule mimics the tetrahedral intermediate during enzymatic catalysis.

**Table 2.** Data collection and refinement statistics.

| **Data collection** | MES (8IBL) | Sulfate (8IBM) |
| --- | --- | --- |
| Wavelength (Å) | 0.9000 | 0.9000 |
| Space group | *P*3 | *P*3 |
| Unit-cell parameters |  |  |
| *a* / *c* (Å) | 83.9 / 64.7 | 83.8 / 64.8 |
| Resolution (Å) | 32-2.60 (2.76-2.60) | 36-2.20 (2.33-2.20) |
| No. of observations | 85,082 | 140,596 |
| No. of unique reflections | 15,681 | 25,685 |
| Completeness (%) | 99.9 (99.8) | 99.6 (98.5) |
| Average *I*/σ(*I*) | 5.8 (1.4) | 10.4 (1.4) |
| Redundancy | 5.4 (5.6) | 5.5 (5.5) |
| *R*<sub>sym</sub> (%) | 27.6 (122.0) | 12.5 (145.5) |
| CC<sub>1/2</sub> (%) | 98.0 (64.2) | 99.7 (74.3) |
| **Refinement** |  |  |
| *R* (%) | 21.0 | 23.0 |
| *R*<sub>free</sub> (%) | 26.4 | 26.6 |
| Average *B* factor (Å<sup>2</sup>) | 37.2 | 48.1 |
| r.m.s. deviation from ideal |  |  |
| Bonds (Å) | 0.002 | 0.002 |
| Angles (°) | 0.55 | 0.47 |
| Ramachandran plot |  |  |
| Favored region (%) | 97.7 | 97.1 |
| Allowed region (%) | 2.3 | 2.9 |
| Outlier region (%) | 0 | 0 |

Values in parentheses are for the highest-resolution shell.

The protein samples were prepared as described previously (*9*). Crystals of the complex of Cut190*SS_S176A and MES were obtained in crystallization buffer containing 0.1 M MES monohydrate pH 6.5, 0.2 M ammonium sulfate, and 30% (w/v) PEG MME 5,000. The protein was concentrated at 7 mg/ml in the buffer containing Tris-HCl pH 8.0, 10 mM NaCl, 5 mM CaCl_2_, and excess molar ratios of synthetic substrate (data not shown). Crystals of the complex of Cut190*SS_S176A and sulfate ion were obtained under the same condition as in the case of the MES complex, without the synthetic substrates. X-ray diffraction experiments were performed at BL-17A at KEK-PF using an automatic data-collection system (*11*), and the data were automatically processed by PReMo (*12*). Manual data processing using XDS (*13*) and truncation using the CCP4 program suite (*14*) were performed as necessary. Phases were determined by the molecular replacement method using PHASER (*15*). Several cycles of manual model rebuilding and refinement were performed by using Coot (*16*) and PHENIX (*17*), respectively.

The structure of the sulfate ion complex (Figure 4c) was determined at 2.2 Å resolution (Table 2). The crystals were soaked into a solution containing 10 mM 2-hydroxyethyl terephthalic acid (MHET) for 30 seconds before data collection. However, no electron density corresponding to MHET was observed at the active site, but large spherical electron density was clearly observed, which is in good agreement with the interpretation of a sulfate ion. Because the crystallization condition contained a high concentration (0.2 M) of ammonium sulfate, we modeled this electron density as a sulfate ion. The bound sulfate in this structure well-matches the sulfo group of MES in the MES-bound structure, which have been accommodated at the oxyanion hole through the formation of hydrogen bonds to the main chain amines of Phe106 and Met177, and the side chain of His254.

Thus, our ligand-binding structures provide structural insight into the molecular mechanism by which the PET molecule is recognized and catalyzed through a tetrahedral intermediate form, although our ligands are slightly different from the original substrate molecule containing an aromatic ring. Based on the above findings, we modeled the PET trimer-bound Cut190 (Figure 4d) as an updated structure from the previously reported binding mode (*19*). This model will be useful for further elucidation of molecular mechanisms of the enzymatic activity based on the intermediate state of substrate binding and cleaving and for the rational design of further-improved enzymes.

## Conclusion

We have demonstrated in this chapter that the enzymatic function of the PET-degrading enzyme Cut190 can be improved by introducing various mutations designed based on the structural information. Both Cut190** and Cut190*SS exhibit higher thermal stability and activity than those of the previously reported mutants. The crystal structure of Cut190**S176A reveals a novel ejecting form in the enzymatic reaction cycle, in which the product would be prompted to dissociate from the active site and substrate exchange is promoted to the next cycle. Structural analysis of the complexes with the ring-shaped ligands, dioxane and MES, along with sulfate provide insights into the aromatic ring-recognition mechanism and also the tetrahedral intermediate during the substrate cleaving. These findings led us to propose a model structure in which the PET trimer is well accommodated in the active site. The proposed model will provide useful structural information for further clarification of elementary processes of enzymatic reaction through computational and other studies.

## Acknowledgements

We sincerely thank Ms. Miho Emori and Ms. Akane Senga for performing the protein purification and crystallization when they were graduate students at Kyoto Prefectural University. The synchrotron radiation experiments were performed with the approval of the Photon Factory Program Advisory Committee (Proposal No. 2019G595). The coordinates and structural data have been deposited in the Protein Data Bank Japan (PDB ID 8IBL and 8IBM for the structures of MES bound and sulfate bound Cut190*SS_S176A, respectively), a member of the Worldwide Protein Data Bank.

## References

1. Oda, M.; Numoto, N.; Bekker, G. J.; Kamiya, N.; Kawai, F., Methods Enzymol 2021, 648, 159–185.

2. Kawai, F.; Oda, M.; Tamashiro, T.; Waku, T.; Tanaka, N.; Yamamoto, M.; Mizushima, H.; Miyakawa, T.; Tanokura, M., Appl Microbiol Biot 2014, 98, 10053–10064.

3. Miyakawa, T.; Mizushima, H.; Ohtsuka, J.; Oda, M.; Kawai, F.; Tanokura, M., Appl Microbiol Biotechnol 2015, 99, 4297–307.

4. Senga, A.; Hantani, Y.; Bekker, G. J.; Kamiya, N.; Kimura, Y.; Kawai, F.; Oda, M., J Biochem 2019, 166, 149–156.

5. Numoto, N.; Kamiya, N.; Bekker, G. J.; Yamagami, Y.; Inaba, S.; Ishii, K.; Uchiyama, S.; Kawai, F.; Ito, N.; Oda, M., Biochemistry 2018, 57, 5289–5300.

6. Oda, M.; Yamagami, Y.; Inaba, S.; Oida, T.; Yamamoto, M.; Kitajima, S.; Kawai, F., Appl Microbiol Biotechnol 2018, 102, 10067–10077.

7. Senga, A.; Numoto, N.; Yamashita, M.; Iida, A.; Ito, N.; Kawai, F.; Oda, M., J Biochem 2021, 169, 207–213.

8. Lu, H.; Diaz, D. J.; Czarnecki, N. J.; Zhu, C.; Kim, W.; Shroff, R.; Acosta, D. J.; Alexander, B. R.; Cole, H. O.; Zhang, Y.; Lynd, N. A.; Ellington, A. D.; Alper, H. S., Nature 2022, 604, 662–667.

9. Emori, M.; Numoto, N.; Senga, A.; Bekker, G. J.; Kamiya, N.; Kobayashi, Y.; Ito, N.; Kawai, F.; Oda, M., Proteins 2021, 89, 502–511.

10. Inaba, S.; Kamiya, N.; Bekker, G. J.; Kawai, F.; Oda, M., J Therm Anal Calorim 2019, 135, 2655–2663.

11. Yamada, Y.; Hiraki, M.; Sasajima, K.; Matsugaki, N.; Igarashi, N.; Amano, Y.; Warizaya, M.; Sakashita, H.; Kikuchi, T.; Mori, T.; Toyoshima, A.; Kishimoto, S.; Wakatsuki, S., AIP Conf Proc 2010, 1234, 415–418.

12. Yamada, Y.; Matsugaki, N.; Chavas, L.; Hiraki, M.; Igarashi, N.; Wakatsuki, S., J Phys Conf Ser 2013, 425.

13. Kabsch, W., Acta Crystallogr D Biol Crystallogr 2010, 66, 125–32.

14. Winn, M. D.; Ballard, C. C.; Cowtan, K. D.; Dodson, E. J.; Emsley, P.; Evans, P. R.; Keegan, R. M.; Krissinel, E. B.; Leslie, A. G.; McCoy, A.; McNicholas, S. J.; Murshudov, G. N.; Pannu, N. S.; Potterton, E. A.; Powell, H. R.; Read, R. J.; Vagin, A.; Wilson, K. S., Acta Crystallogr D Biol Crystallogr 2011, 67, 235–42.

15. McCoy, A. J.; Grosse-Kunstleve, R. W.; Adams, P. D.; Winn, M. D.; Storoni, L. C.; Read, R. J., J Appl Crystallogr 2007, 40, 658–674.

16. Emsley, P.; Lohkamp, B.; Scott, W. G.; Cowtan, K., Acta Crystallogr D Biol Crystallogr 2010, 66, 486–501.

17. Adams, P. D.; Afonine, P. V.; Bunkoczi, G.; Chen, V. B.; Davis, I. W.; Echols, N.; Headd, J. J.; Hung, L. W.; Kapral, G. J.; Grosse-Kunstleve, R. W.; McCoy, A. J.; Moriarty, N. W.; Oeffner, R.; Read, R. J.; Richardson, D. C.; Richardson, J. S.; Terwilliger, T. C.; Zwart, P. H., Acta Crystallogr D Biol Crystallogr 2010, 66, 213–21.

18. Tournier, V.; Topham, C. M.; Gilles, A.; David, B.; Folgoas, C.; Moya-Leclair, E.; Kamionka, E.; Desrousseaux, M. L.; Texier, H.; Gavalda, S.; Cot, M.; Guemard, E.; Dalibey, M.; Nomme, J.; Cioci, G.; Barbe, S.; Chateau, M.; Andre, I.; Duquesne, S.; Marty, A., Nature 2020, 580, 216–219.

19. Kawabata, T.; Oda, M.; Kawai, F., J Biosci Bioeng 2017, 124, 28–35.

